# Experimental safety testing shows that the NSAID tolfenamic acid is not toxic to *Gyps* vultures in India at concentrations likely to be encountered in cattle carcasses

**DOI:** 10.1101/2021.08.23.456758

**Authors:** S Chandramohan, John W. Mallord, Kesavan Manickam, A. K. Sharma, K. Mahendran, Reena Gupta, Krishna Chutia, Karikalan Mathes, Abhijit Pawde, Nikita V. Prakash, P. Ravichandran, Debasish Saikia, Rohan Shringarpure, Avinash Timung, Toby H. Galligan, Rhys E. Green, Vibhu M. Prakash

## Abstract

Population declines of *Gyps* vultures across the Indian subcontinent were caused by unintentional poisoning by the non-steroidal anti-inflammatory drug (NSAID) diclofenac. Subsequently, a number of other NSAIDs have been identified as toxic to vultures, while one, meloxicam, is safe at concentrations likely to be encountered by vultures in the wild. Other vulture-safe drugs need to be identified to reduce the use of those toxic to vultures. We report on safety-testing experiments on the NSAID tolfenamic acid on captive vultures of three *Gyps* species, all of which are susceptible to diclofenac poisoning. Firstly, we estimated the maximum level of exposure (MLE) of wild vultures and gave this dose to 38 Near Threatened *G. himalayensis* by oral gavage, with 15 control birds dosed with benzyl alcohol (the carrier solution for tolfenamic acid). Two birds given tolfenamic acid died with elevated uric acid levels and severe visceral gout, while the remainder showed no adverse clinical or biochemical signs. Secondly, four *G. himalayensis* were fed tissues from water buffaloes which had been treated with double the recommended veterinary dose of tolfenamic acid prior to death and compared to two birds fed uncontaminated tissue; none suffered any clinical effects. Finally, two captive Critically Endangered vultures, one *G. bengalensis* and one *G. Indicus*, were given the MLE dose by gavage and compared to two control birds; again, none suffered any clinical effects. The death of two *G. himalayensis* may have been an anomaly due to i) the high dose level used and ii) the high ambient temperatures at the time of the experiment. Tolfenamic acid is likely to be safe to *Gyps* vultures at concentrations encountered by wild birds and could therefore be promoted as a safe alternative to toxic NSAIDs. It is manufactured in the region, and is increasingly being used to treat livestock.

## 1. Introduction

Three species of *Gyps* vultures endemic to southern Asia, White-rumped *G. bengalensis*, Indian *G. indicus* and Slender-billed *G. tenuirostris* Vultures, underwent population declines in the Indian subcontinent, beginning in the mid-1990s (Prakash *et al.* 2007, Chaudhary *et al.* 2012). In India, the population of *G. bengalensis* had declined by 99.9% by 2007, relative to the early 1990s, while that of *G. indicus* and *G. tenuriostris* combined declined by 96.8% (Prakash *et al.* 2007). All three species are classified as Critically Endangered (Birdlife International 2021). The cause of these declines was identified as unintentional poisoning by the veterinary non-steroidal anti-inflammatory drug (NSAID) diclofenac (Oaks *et al.* 2004, Shultz *et al.* 2004, Green *et al.* 2004), which vultures were exposed to when they fed on carcasses from domesticated ungulates that had been treated with the drug within a few days before death. Vultures died from kidney failure, with macroscopic signs at post-mortem examination being extensive visceral gout (formation of uric acid crystals within and coating tissues, especially kidney and liver) (Oaks *et al.* 2004). Histopathological examination of kidney tissue of *G. bengalensis* exposed to diclofenac revealed severe necrosis of the convoluted tubules and large aggregates of urate crystals (Meteyer *et al.* 2005). The concentration of uric acid in the blood serum of captive *G. bengalensis*, African white-backed *G. africanus* and Eurasian griffon *G. fulvus* vultures dosed experimentally with diclofenac increased markedly and to a similar extent across species, often to ten times the normal level, within 24h of treatment. Experimentally-treated birds died within a few days (Oaks *et al.* 2004; Swan *et al.* 2006a). As a result of the overwhelming evidence implicating diclofenac in the vulture declines, the governments of India, Pakistan and Nepal banned the manufacture, importation and veterinary use of diclofenac in 2006, with Bangladesh doing so in 2010. The ban has contributed to slowing or halting declines in India and Pakistan (Chaudhry *et al.* 2012, Prakash *et al.* 2017), and enabled partial recovery in Nepal (Galligan *et al.* 2019).

Diclofenac, which is used legally by humans, remains on sale for veterinary use illegally in India despite its ban (Galligan *et al.* 2020). In addition, several other NSAIDs are available for approved veterinary use in pharmacies across the subcontinent (Galligan *et al.* 2020). Of these, experimental safety testing on captive vultures has shown that ketoprofen and carprofen are toxic to vultures (Naidoo *et al.* 2010; Fourie *et al.* 2015; Naidoo *et al.* 2018). Aceclofenac is a pro-drug of diclofenac and is rapidly metabolized into diclofenac in cattle, so will have the same disastrous effects on vultures (Galligan *et al.* 2016). In addition, wild *Gyps* vultures have been recovered dead in Spain *(G. fulvus)* with co-occurrence of extensive visceral gout with tissue residues of flunixin (Zorilla *et al.* 2014, Herrero-Villar *et al.* 2020) and in India (*G. bengalensis)* with residues of nimesulide (Cuthbert *et al.* 2016; Nambirajan et al. 2021), which we take to be strong evidence that these two drugs are also nephrotoxic to vultures.

Currently, the only NSAID that has been shown through experimental safety testing not to be toxic to vultures is meloxicam, which was found not to cause death or morbidity in captive *Gyps* vultures, nor elevate the concentration of uric acid in their blood serum, even at doses higher than the likely maximum level of exposure of wild birds (Swan *et al.* 2006b, Swarup *et al.* 2007). As a result, this drug has been promoted throughout South Asia as an alternative to diclofenac, has become more commonly offered for use on livestock, especially in Nepal (Galligan *et al.* 2020), and has increased in prevalence in surveys of NSAID levels in cattle and water buffalo carcasses available to vultures in India (Cuthbert *et al.* 2014). As expected from the experimental studies, there was no indication that two wild *G. bengalensis* found dead in India with meloxicam residues in the liver had died because of nephrotoxic effects of the drug (Cuthbert et al. 2016). In addition, seven *G. fulvus* found dead with meloxicam residues in the liver in Spain, where the drug is also used for veterinary purposes, had died for reasons unrelated to nephrotoxicity (Herrero-Villar *et al.* 2020). Although it remains important to identify and ban veterinary use of other NSAIDs which are nephrotoxic to vultures, as proposed in the Indian government’s *Vulture Action Plan 2020-2025* (MoEFCC 2020), it is also a high priority to find vulture-safe NSAIDs in addition to meloxicam to give veterinarians and livestock owners more choice.

Tolfenamic acid is used in veterinary medicine because of its anti-inflammatory, antipyretic and analgesic properties (Sidhu *et al.* 2010). It is also effective, in conjunction with antibiotics, for the treatment of respiratory disease and mastitis in cattle (Delaforge *et al.* 1994, EMEA 1997, Patel *et al.* 2018). The normal therapeutic dose in cattle is 2 mg kg^-1^ body weight (b.w.) day^-1^ for two days, or a single injection of 4 mg kg^-1^ b.w. Undercover pharmacy surveys in the subcontinent, especially in Nepal and Bangladesh, reveal that tolfenamic acid is increasingly being offered for sale for veterinary use, although still at a lower frequency than several other NSAIDs (Galligan *et al.* 2020). In this paper, we report results from experiments on the safety of tolfenamic acid to captive *Gyps* vultures. We estimated a precautionary maximum level of exposure (MLE) of tolfenamic acid to wild vultures and then administered this dose to captive vultures by gavage. We also dosed water buffaloes with twice the recommended dose of the drug prior to slaughter and fed their tissues to captive vultures.

## 2. Methods

### 2.1 Trial animals, housing and management

The experiments were carried out at the Vulture Conservation Breeding Centre at Pinjore, Haryana, India in separate phases in November 2017, February 2018, and April, July and November 2019. Phases I-V of the safety testing were carried out on captured wild Himalayan Griffon Vultures (hereafter, HG), which are regular winter visitors to the centre at Pinjore from breeding populations further north in Asia. Wild HGs were attracted by seeing captive vultures in aviaries and by a supplementary feeding station, and were trapped in a large (27 × 5 × 9 m) baited walk-in cage trap. They were transferred to a four-compartment purpose-built aviary, each compartment being 6 × 6 × 5 m, and holding a maximum of four vultures. Twenty HGs intended for use in Phases I-III were trapped on 6^th^ February 2017, but seven birds died during an outbreak of Avian TB, leaving 13 birds for testing. A health check was performed on these birds on 23^rd^ October before the experiments were carried out on 6^th^ and 28^th^ November 2017 and 15^th^ February 2018. All birds were released on 30^th^ April 2018, 14 months and 24 days after initial capture. HGs (n = 42) for Phases IV and V were trapped on 16^th^ March 2019, with the health check done a week later and the experiments carried out between 7^th^ April and 26^th^ May. These birds were released on 29^th^ September 2019, six months and 29 days after initial capture. Birds were fed an average of 2.5 kg of goat meat, sourced to be free of NSAIDs, twice per week, and received routine health monitoring, throughout their time in captivity. In Phase VI (28^th^ November 2019), testing was carried out on one captive *Gyps bengalensis* and one *G. indicus*, with one more bird of each species serving as controls. These birds were part of the captive population, but were not suitable for conservation breeding or release because of injuries in the wild prior to capture. A health check was performed prior to the experiment and, apart from these injuries, the birds were confirmed to be in good health. These birds were housed individually in a 6 × 6 × 5 m aviaries, to facilitate the provision of food, and maintained on their normal feeding regime, i.e., 2 kg of goat meat per bird twice a week. Prior to the experiment, birds were fed solely on goat liver to acclimate them to this source of food.

### 2.2 Treatment and study design for oral gavage experiments

Phases I-III each involved treatment and control groups, with HGs being assigned at random to a tolfenamic acid (TA)-treated group (*n* = 2, 2 & 5, respectively in Phases I-III respectively) or a control group (*n* = 2, 2 & 4, respectively). Larger numbers of HGs were assigned to the Phase IV.1 trial (treatment n = 12, control n = 3) and the Phase IV.2 trial (treatment n = 17, control n = 4; Table 1). Of the 49 HGs used in Phases I-V, 32 birds were used only once in a TA-treatment group and 10 birds were used once in a control group. Seven birds were included in more than one Phase (treatment / treatment = 3; control / control = 3; control / control / treatment = 1), with a gap of at least three months between consecutive experiments. Experiments on the same individual were a minimum of 11 weeks apart for birds given sham treatments and over a year apart for birds dosed at least once with tolfenamic acid. The dose rate was set at a level estimated to be a precautionary maximum level of exposure (MLE) to tolfenamic acid of wild vultures from carcasses of ungulates treated with the drug. The MLE was calculated by first estimating a value for the concentration of tolfenamic acid near to the upper limit expected in the livers of cattle treated with the drug (Supplementary Material: Appendix 1). The dose of tolfenamic acid required was then calculated for each bird as the quantity of drug expected to be ingested by the bird if it ate a meal large enough to provide its energetic requirements for three days. The size of this meal was calculated from the bird’s body weight. The calculated doses were 3.2-3.7 mg kg^-1^ vulture body weight (v.b.w.). Control HGs were sham-treated by gavage with benzyl alcohol (the carrier solution for TA). In Phase VI, one *G. bengalenesis* and one *G. indicus* were dosed by oral gavage with MLE of 4.0-4.5 mg kg^-1^ v.b.w. Control birds (one of each species) were sham-treated by gavage with benzyl alcohol. Birds were observed following dosing for any regurgitation, but none occurred. Vultures were observed at regular intervals for the following seven days for any signs of ill-health.

**Table 1.** Schedule and results of experimental safety testing of tolfenamic acid on Himalayan Griffon *Gyps himalayensis* (HG),White-rumped *G. bengalensis* (WRV) and Indian *G. indicus* (IV) Vultures, and the number of deaths recorded in each Phase. No control birds died.

| Date | Species | Phase | Mean body weight (kg) | Mean dose (mg kg<sup>-1</sup> v.b.w.) | Route | N dosed | N died | % mortality | N Control |
| --- | --- | --- | --- | --- | --- | --- | --- | --- | --- |
| 6 Nov 2017 | HG | I | 9.1 | 3.289 | Gavage | 2 | 0 | 0 | 2 |
| 27 Nov 2017 | HG | II | 8.75 | 3.339 | Gavage | 2 | 0 | 0 | 2 |
| Feb 2018 | HG | III | 8.72 | 3.345 | Gavage | 5 | 0 | 0 | 4 |
| Apr 2019 | HG | IV.1 | 7.7 | 3.507 | Gavage | 12 | 2 | 16.7 | 3 |
| July 2019 | HG | IV.2 | 8.3 | 3.408 | Gavage | 17 | 0 | 0 | 4 |
| July 2019 | HG | V |  | 0.072-0.112 | Tissue | 4 | 0 | 0 | 2 |
| Nov 2019 | WRV / LBV | VI |  | 4.0-4.5 | Gavage | 2 | 0 | 0 | 2 |
| **Totals** |  |  |  |  |  | **44** | **2** | **4.5** | **19** |

A commercial brand of veterinary tolfenamic acid was used (Maxxtol, Intas Pharmaceuticals, Ahmedabad, India), which is widely available and was purchased from several pharmacies near Pinjore. The product had a stated tolfenamic acid concentration of 40 mg l^-1^.

### 2.3 Phase V treatment and design

In Phase V, vultures were fed meat from domesticated water buffaloes *Bubalus bubalis* that had been treated with tolfenamic acid. Three buffalo steers of about 11 months of age and weighing approximately 220 kg received a single intramuscular injection of tolfenamic acid at a dose of 4 mg kg^-1^ b.w. in the neck. This is twice the recommended veterinary dose. We used a high does because cattle and buffaloes in India are often treated with doses of NSAIDs much higher than the recommended dose, as is apparent from measured concentrations in the livers of dead animals sampled at carcass dumps (Green *et al.* 2007). Two of the animals were slaughtered 24 hours after treatment. The third animal, which was not treated with tolfenamic acid, was also slaughtered to provide meat for vultures in the control group. Buffaloes were euthanized with a saturated solution of MgSO_4_ at 2 ml kg^-1^b.w. Samples were collected from the injection site (i.e., neck muscle), liver and kidney to measure tolfenamic acid concentrations. The concentrations of tolfenamic acid in the kidneys of two buffaloes 24 hours after treatment were 0.800 and 1.250 mg kg^-1^ wet weight (w.w.). Concentrations were 0.596 and 0.399 mg kg^-1^ (w.w.) for liver and 0.027 and 0.053 mg kg^-1^ (w.w.) for muscle. Four HGs were fed the maximum likely meal, i.e. enough to satisfy the birds’ energy requirements for three days, 1.280-1.428 kg of liver tissue from the treated buffalo; this is equivalent to a dose of tolfenamic acid 0.072-0.112 mg kg^-1^ v.b.w. Two control HGs were given similar quantities of meat from the untreated buffalo. Birds were kept under observation to ensure that all the meat was eaten and not regurgitated. Vultures were observed at regular intervals for the following seven days for any signs of ill-health.

### 2.4 Blood sampling

In all phases of the study, blood samples were collected by direct veno-puncture from the brachial or tarsal veins. A total of approximately 15 ml of blood (about 3% of estimated blood volume) was collected from each vulture over a 7-d period. In Phases I-IV and VI, blood serum taken from each bird at 0 (i.e. pre-treatment), 2, 6, 12, 24, 36, 48, 96 and 168 hours after treatment were analysed to estimate the concentration of the following: uric acid, creatinine, urea, alanine aminotransferase (ALT), total protein, albumin, calcium, phosphorous and chloride. In Phase V, samples were taken from each bird at 0 (i.e., pre-treatment), 48 and 168 hours after the birds ate the liver tissue for analysis of the same set of blood parameters.

### 2.5 Extraction and measurement of tolfenamic acid in plasma and tissues

Tolfenamic acid was extracted from vulture plasma and tissue samples using protein precipitation method, and was quantified using liquid chromatography tandem mass spectrometry (LC-MS/MS) with Electro Spray Ionization (ESI) and multiple reaction monitoring (MRM) in negative ionization mode.

Tolfenamic acid was extracted from a 0.5-0.58 g samples of kidney, liver and injection site (neck muscle) tissues from two dosed buffaloes. The tissues were homogenised with 2 ml of HPLC grade acetonitrile, which was then centrifuged at 6000 rpm for 6 min. After the centrifugation was complete, the supernatant was decanted into 2 ml vials through a PTFE filter. The vials were stored at -20°C before analysing them with LC-MS/MS.

Dimethylsulfoxide (DMSO, 1.16 mL) was added to 1.16 mg of tolfenamic acid in a 1.5 mL micro-centrifuge tube, mixed well and sonicated; 1 mL DMSO was added to 5 mg of tolfenamic acid (D-4) and used as internal standard stock. Analyte working calibration standards (32 to 20000 ng mL^-1^) and quality control samples (QC) of tolfenamic acid were prepared in DMSO. An internal working standard solution was prepared by diluting 0.010 mL of the internal standard stock solution to 100 mL with acetonitrile to provide a concentration of 1.0 μg mL^-1^. This solution was mixed well and stored at 2 to 8°C. Calibration standards and quality control samples of tolfenamic acid were prepared by spiking with 2.5 μL of the analyte working solution in 47 μL of blank vulture plasma. Study samples were aliquoted into Eppendorf tubes and 10 μL of the internal working standard solution (1.50 μL ml^-1^ of D4-IS) added. Samples were quenched with 250 μL of acetonitrile, vortexed and centrifuged at 14,000 rpm for 10 minutes at 4°C. 150 μL of supernatant was transferred to 1 mL vials for analysis in LC-MS/MS.

### 2.6 Measurement of serum constituents

Biochemical parameters were analyzed using commercial kits (Coral Clinical Systems, Tulip Diagnostics) with GENESYS 10UV spectrophotometer (Thermo Scientific). Total protein was estimated using the Biuret method and ALT the Reitman and Frankel method. Creatinine, urea and uric acid concentrations were quantitated using immuno-inhibition / modified IFCC method, alkaline picrate method, DAM method and uricase/PAP method respectively. Serum calcium and phosphorus concentrations were estimated with OCPC and modified Gomorri’s method respectively, and chloride with thiocyanate.

### 2.7 Statistical analysis

We calculated Clopper-Pearson exact binomial 95% confidence limits (Clopper & Pearson 1934) for the proportion of birds that died in our experiments. We used a two-tailed Fisher exact test to assess the statistical significance of the difference in the proportion of HG that died during the experiment between treatment and control groups for pooled data from all experiments in which tolfenamic acid was administered to HG by oral gavage (Phases I – IV.2). We also used two-tailed Fisher exact tests to compare the proportion of HG treated with tolfenamic acid by gavage that died with the proportions of *Gyps* vultures that died when experimentally administered diclofenac and meloxicam by gavage in other experiments published elsewhere. These pooled data were from *G. bengalensis, G. africanus* and *G. fulvus* treated with diclofenac by gavage in experiments reported by Oaks *et al.* (2004), Swan *et al.* (2006a) and Swan *et al.* (2006b). For meloxicam, we used results for *G. africanus, G. bengalensis* and *G. indicus* given a meloxicam dose greater than the MLE by gavage from Swan *et al.* (2006).

In Phases I-V, the effect of tolfenamic acid on changes in the concentration of the various blood parameters was analysed by comparing treated with control birds using a *t-*test. The dependent variable was the ratio of the mean serum concentration at each time interval to the serum concentration immediately pre-treatment (i.e., 0 hours). Ratios greater than one would indicate an increase in concentration as a result of the treatment.

### 2.8 Animal ethics

Permission to dose and slaughter buffaloes in Phase V experiments was obtained from the Committee for the Purpose of Control and Supervision of Experiments on Animals (CPCSEA, No.F. 26-1/2015-16/JD(R)). Permission to carry out experiments on vultures was granted by the Government of Haryana Forest Department, and the RSPB’s Animal Ethics Committee (EAC2016-02).

## 3. Results

### 3.1 Safety testing on HG by administration of tolfenamic acid by oral gavage (Phases I-IV)

The mean concentration of tolfenamic acid in the blood serum of treated vultures was highest two hours after the birds were dosed, and declined steadily thereafter (SOM Appendix 2: Table S2). Two of the 38 HG (5.3%; 95% confidence limits 0.6 – 17.8%) treated with tolfenamic acid during Phases I to IV.2 in April 2019 died 36-48 hours post-treatment (Table 1). Both birds were in the treatment group of Phase IV.1 in April 2019. All other treated birds and controls in these experiments survived with no apparent ill-effects. The difference between the proportion of birds that died was not significantly different between the treatment and control groups (Fisher exact test, P = 1.000).

The mean uric acid concentration in the blood plasma was higher in treatment group than the control group at all sampling times, including before treatment, but the difference was only significant at 2h and 6h after treatment (Fig. 1, Table 2). When the serum uric acid concentration at each sampling time after treatment was expressed as a ratio relative to the pre-treatment value for that bird, there were no significant differences between treatment and control groups for any sampling time and no indication of a general tendency for mean before / after uric acid ratio to differ consistently between treatment and control groups (Table 2). Mean blood serum alanine aminotransferase (ALT) concentrations also showed no consistent tendency to differ between treatment and control groups (Table 3), though it was significantly lower in treated birds than in controls at one sampling period (2h: Table 3).

**Figure 1.**
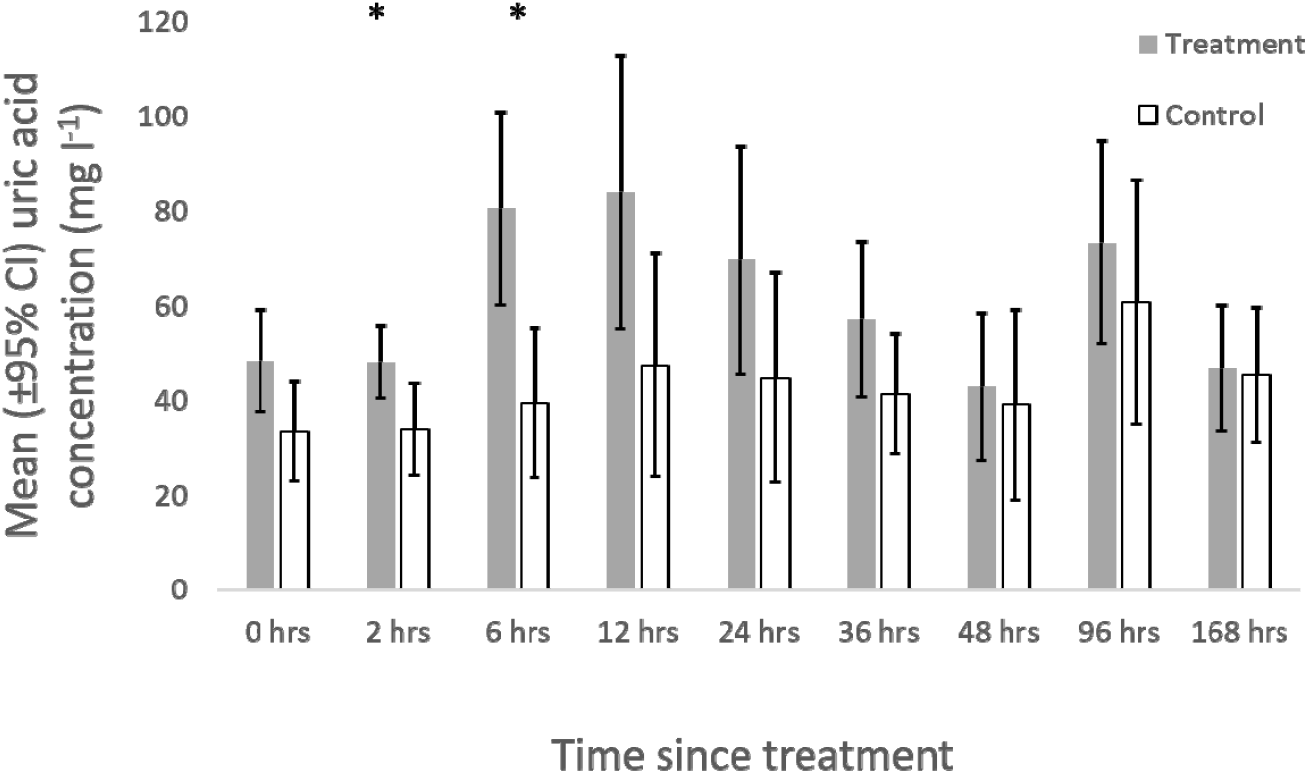
Effect of administration of tolfenamic acid by oral gavage on the concentration of uric acid in the blood serum of Himalayan Griffon Vultures *Gyps himalayensis*. Asterisks denote a significant difference between treatment and control groups.

**Table 2.** The effect of tolfenamic acid treatment on blood serum uric acid levels, expressed as a) uric acid concentrations, and b) the ratio between post- and pre-treatment (i.e. 0 hours) uric acid concentrations in the blood serum of Himalayan Griffon Vultures *Gyps himalayensis* dosed by oral gavage. In b), values > 1 indicate an increase in concentrations post-treatment.

| a) Mean uric acid concentration<br>(mg l<sup>-1</sup>) |  |  |  |  | b) Mean ratio of post-:pre-treatment<br>levels |  |  |  |
| --- | --- | --- | --- | --- | --- | --- | --- | --- |
| Time h | Control (n) | Dosed (n) | *t* | *p* | Control | Dosed | *t* | *p* |
| 0 | 33.5 (15) | 48.3 (36) | 1.66 | 0.10 |  |  |  |  |
| **2** | **33.9 (15)** | **48.0 (36)** | **2.39** | **0.04** | 1.09 | 1.10 | 0.17 | 0.87 |
| **6** | **39.4 (15)** | **80.5 (36)** | **2.52** | **0.02** | 1.59 | 1.78 | 1.86 | 0.068 |
| 12 | 47.3 (15) | 83.9 (36) | 1.57 | 0.12 | 1.94 | 1.99 | 0.57 | 0.57 |
| 24 | 44.8 (15) | 69.7 (36) | 1.27 | 0.21 | 1.48 | 1.55 | 0.19 | 0.85 |
| 36 | 41.2 (13) | 57.1 (34) | 1.17 | 0.25 | 1.41 | 1.19 | 0.93 | 0.36 |
| 48 | 39.0 (15) | 42.8 (36) | 0.28 | 0.78 | **1.25** | **0.91** | **2.14** | **0.037** |
| 96 | 60.8 (11) | 73.2 (19) | 0.77 | 0.45 | 2.57 | 2.07 | 1.23 | 0.23 |
| 168 | 45.5 (8) | 46.8 (9) | 0.15 | 0.88 | 2.28 | 1.96 | 0.69 | 0.50 |

**Table 3.** Concentration of blood serum alanine aminotransferase (ALT) after treatment of Himalayan Griffon Vultures *Gyps himalayensis* by oral gavage with tolfenamic acid.

| Time h | Mean ALT concentration(Units l<sup>-1</sup>) |  | *t* | *p* |
| --- | --- | --- | --- | --- |
|  | Control (n) | Dosed (n) |  |  |
| 0 | 34.3 (15) | 36.0 (36) | 0.35 | 0.73 |
| **2** | **30.8 (15)** | **22.8 (36)** | **2.11** | **0.04** |
| **6** | 37.5 (15) | 37.9 (36) | 0.06 | 0.96 |
| 12 | 36.6 (15) | 40.0 (36) | 0.49 | 0.63 |
| 24 | 37.8 (15) | 43.4 (36) | 1.32 | 0.19 |
| 36 | 40.8 (15) | 42.6 (36) | 0.44 | 0.66 |
| 48 | 44.0 (15) | 43.2 (36) | 0.13 | 0.89 |
| 96 | 23.4 (11) | 26.9 (19) | 0.60 | 0.55 |
| 168 | - | - | - | - |

There was no indication of a difference in the mean concentration of tolfenamic acid in the blood serum of treated birds that survived and the two birds that died at any sampling time (Table S2). However, the two birds that died had a marked elevation of the mean before / after serum uric acid concentration compared with the treated birds that survived and the controls from 6h after treatment until death (Fig. 2). The highest before / after ratio in the surviving treated birds was 2.0, compared to an average 27-fold increase after 36 hours in the two birds that died. There were also few significant differences between treated and control birds in the change in concentration of the other biochemical parameters measured (Appendix 2, Table S2).

**Figure 2.**
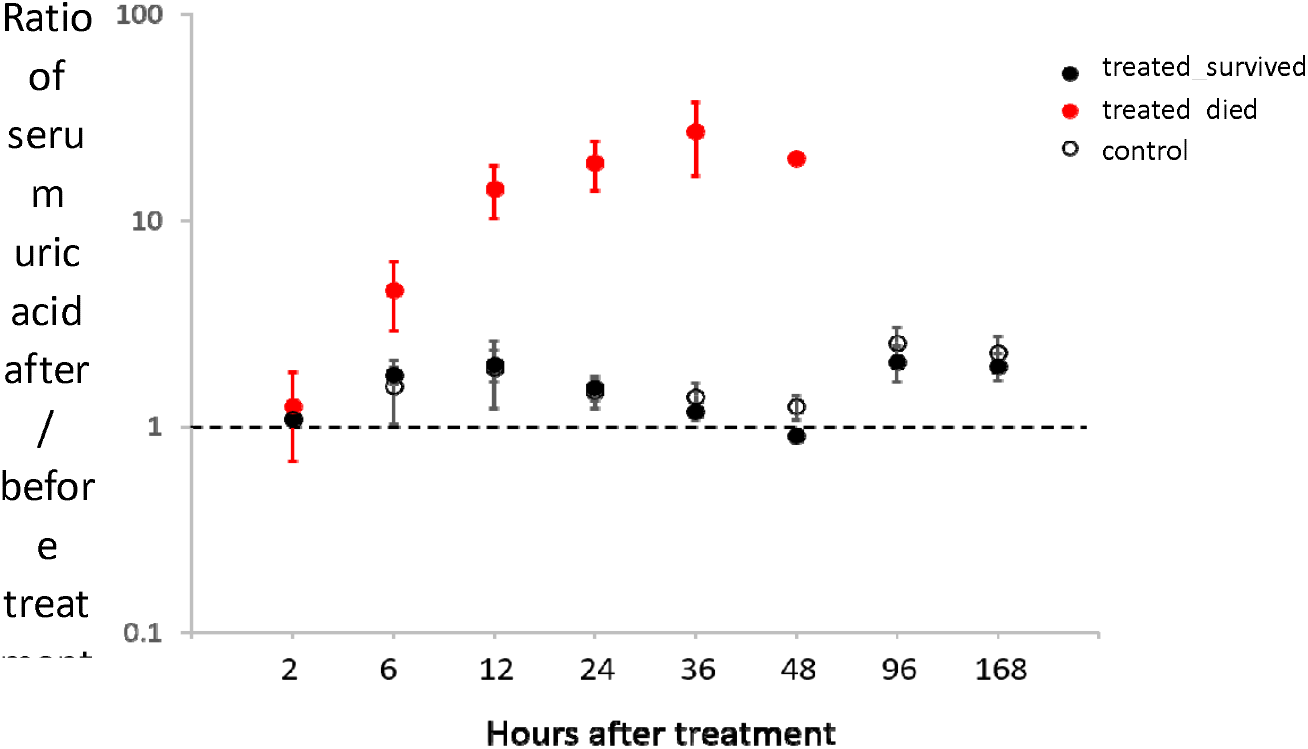
Effect of administration of tolfenamic acid by oral gavage on uric acid in the serum of Himalayan Griffon Vultures *Gyps himalayensis*. Symbols show the ratio of the mean serum concentration (± SE) of uric acid after treatment to that immediately before treatment for birds that were a) treated with tolfenamic acid and survived (n = 36), b) treated and died (n = 2) and c) controls(n = 15) that were dosed by oral gavage with benzyl alcohol. Values > 1 indicate an increase in uric acid concentration post-treatment. Increase in uric acid in both treatment and control groups at 96 hours followed feeding.

### 3.2. Post-mortem analysis of dead G. himalayensis

Post-mortem analysis was carried out on the two *G. himalayensis* that died following oral gavage with tolfenamic acid. Both birds were dosed on 8^th^ April 2019; the first began to appear unwell on the evening of the 9^th^, and died early in the morning of the 10^th^, while the other died on the 11^th^.

Formalin-fixed tissue samples from the liver, lungs, kidney, spleen, heart, brain, ureter, stomach and intestine of both birds were examined. Both birds showed similar pathologies: the kidneys showed severe damage of tubules characterised by moderate to severe tubular dilation and necrosis with radiated needle-like urate crystals in the lumen. Kidney histological changes were more extensive with complete destruction of the renal tubules with swollen, degenerated and putrescent epithelial cells. The interstitial blood vessels were severely engorged with mononuclear cell infiltration. Similar lesions were also observed in the pericardium of the heart, serosal surface of the lung, spleen and liver. The diagnosis for both birds was visceral gout resulting in multi-organ pathology.

### 3.3 Comparison of death rate between HG treated with tolfenamic acid and other Gyps species treated with diclofenac and meloxicam

Of the HG to which tolfenamic acid was administered by gavage, 2 birds of 38 died, compared with eight out of nine *Gyps* vultures treated with diclofenac by gavage and none of 45 *Gyps* vultures treated with meloxicam by gavage. Hence, the lower proportions of deaths in experiments with tolfenamic acid and meloxicam than with diclofenac were each highly statistically significant (Fisher exact test, both P < 0.0001). The proportion of deaths in the tolfenamic acid experiment was not significantly different from that with meloxicam (Fisher exact test, P = 0.207).

### 3.4 Safety testing by feeding HG on tissues of tolfenamic acid-treated water buffaloes (Phase V)

All of the HG fed on tissues of tolfenamic acid-treated water buffaloes survived beyond the end of the experiment, as did the controls (Table 1). Detectable concentrations of tolfenamic acid in the blood serum of treated vultures were observed only 48 hrs after the birds were dosed (SOM Appendix 2: Table S3). Tolfenamic acid detected in a single bird in the control group was presumably due to contamination. There were no significant differences in mean before / after uric acid blood serum ratio between treatment and control groups for any sampling time, the largest difference being at 48h post-treatment (treatment mean = 1.456; control mean = 0.803; *t* = 2.01, *P* = 0.11, Table 4). Mean blood serum alanine aminotransferase (ALT) concentrations also showed no tendency to differ between treatment and control groups (Table 5). There were also few significant differences between treated and control birds in the change in concentration of the other biochemical parameters measured (Appendix 2, Table S3).

**Table 4.** The effect of tolfenamic acid treatment on blood serum uric acid levels, expressed as a) uric acid concentrations, and b) the ratio between post- and pre-treatment (i.e., 0 hours) uric acid concentrations in the blood serum of Himalayan Griffon Vultures *Gyps himalayensis* fed contaminated buffalo tissue. In b), values > 1 indicate an increase in concentrations post-treatment.

| a) Mean uric acid concentration<br>(mg l<sup>-1</sup>) |  |  |  |  | b) Mean ratio of post-:pre-treatment<br>levels |  |  |  |
| --- | --- | --- | --- | --- | --- | --- | --- | --- |
| Time h | Control<br>(n = 2) | Dosed<br>(n = 4) | *t* | *p* | Control | Dosed | *t* | *p* |
| 0 | 65.9 | 63.5 | 0.33 | 0.76 |  |  |  |  |
| 48 | 49.4 | 90.2 | 1.42 | 0.23 | 0.80 | 1.46 | 2.01 | 0.11 |
| 168 | 52.2 | 40.4 | 1.87 | 0.13 | 0.84 | 0.71 | 0.50 | 0.64 |

**Table 5.**
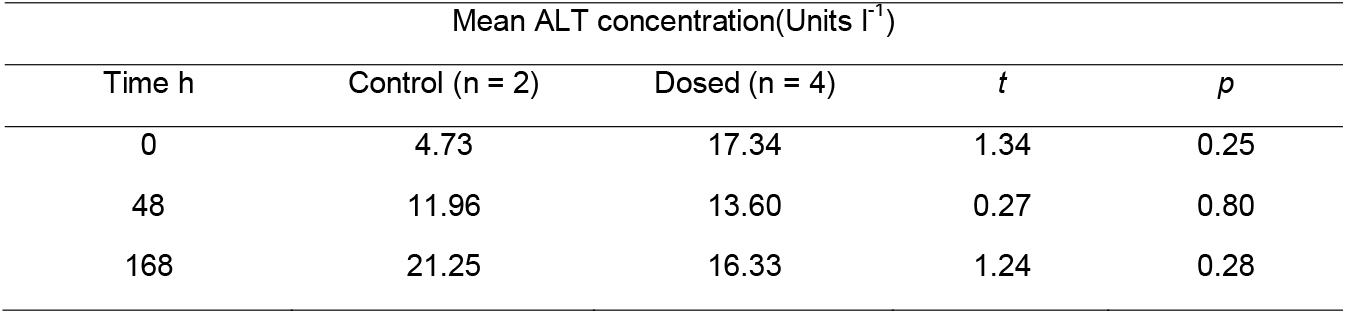
Concentration of blood serum alanine aminotransferase (ALT) after treatment of Himalayan Griffon Vultures *Gyps himalayensis* with tolfenamic acid, after feeding on contaminated buffalo tissue.

### 3.5 Safety testing on G. bengalensis and G. indicus by administration of tolfenamic acid by oral gavage (Phase VI)

Neither individual treated with tolfenamic acid showed any ill effects and both survived until the end of the experiment, 168 hours post-treatment (Table 1). These birds remained in captivity and are both alive more than one year after the experiment. The two control birds also survived and showed no ill-effects. There was no indication of any consistent elevation of mean blood serum uric acid concentration at any sampling time (Table 6), though the small samples preclude any statistical testing.

**Table 6.** The effect of tolfenamic acid treatment on mean blood serum uric acid concentration expressed as a) uric acid concentrations, and b) the ratio between post- and pre-treatment (i.e. 0 hours) uric acid concentrations in the serum of White-rumped *Gyps bengalensis* and Indian *G.indicus* Vultures. In b), values > 1 indicate an increase in concentrations post-treatment.

| Time<br>h | a) Mean uric acid<br>concentrations |  | b) Mean ratio of<br>post:pre-treatment<br>levels |  |
| --- | --- | --- | --- | --- |
|  | Control<br>(n = 2) | Dosed<br>(n = 2) | Control | Dosed |
| 0 | 4.13 | 3.66 |  |  |
| 2 | 4.95 | 5.39 | 1.21 | 1.44 |
| 6 | 3.78 | 4.28 | 0.92 | 1.16 |
| 12 | 5.49 | 6.13 | 1.33 | 1.65 |
| 24 | 4.49 | 5.54 | 1.10 | 1.49 |
| 36 | 5.21 | 4.87 | 1.26 | 1.32 |
| 48 | 3.68 | 3.42 | 0.90 | 0.93 |
| 96 | 5.09 | 4.13 | 1.25 | 1.12 |
| 168 | 5.65 | 5.38 | 1.38 | 1.45 |

## 4. Discussion

The results of our study indicate that, although it was not completely safe at the dose level administered, veterinary use of tolfenamic acid poses a much lower risk to wild vultures than does diclofenac, the drug responsible for the catastrophic declines of Asian vulture populations from the mid-1990s. However, two Himalayan Griffon Vultures died after treatment with tolfenamic acid, whereas no deaths caused by the established vulture-safe drug meloxicam have been documented, either in experiments on captive birds or in wild vultures. For this reason, some caution and careful consideration of the deaths of these two birds are needed.

A large majority (95%) of Himalayan Griffons that were given tolfenamic acid by oral gavage survived. Both birds died during the same experiment (Phase IV.1). Although the group of birds used in this experiment was trapped just three weeks before the beginning of the experiment, we think it unlikely that the stress of being taken into captivity itself caused the birds’ deaths. Both of them showed elevated uric acid levels typical of NSAID poisoning (Oaks *et al.* 2004, Swan *et al.* 2006, Naidoo *et al.* 2010), whereas uric acid levels tend to decrease in response to most types of stress (Gormally *et al.* 2018, 2019). However, it remains possible that stress affected the birds’ sensitivity to tolfenamic acid. This experiment was conducted in April 2019, shortly before the onset of the spring monsoon rains, when temperatures reached 40°C. Although no studies have looked at the impact of heat stress on uric acid metabolism in birds, in humans it can cause both rises in uric acid levels and kidney failure (Roncal-Jimenez *et al*.2015, Wesseling *et al.* 2016).

In addition to speculating that stress caused by capture and high temperatures may have been contributory causes of the deaths of these birds, we also note that the doses of tolfenamic acid given to all birds may have been substantially higher than those in most meals of wild vultures feeding on a contaminated cow carcass are likely to be. This is because of the precautionary approach we adopted in calculating the MLE concentration in vulture food and its sensitivity to the arbitrary choices we made (see SOM Appendix 1). Our approach is precautionary in the following three ways. (1) We used that ratio of the 95^th^ percentile of the distribution of diclofenac levels in cow carcass samples, which is a high and arbitrary value. (2) We used the ratio of that carcass survey to the mean diclofenac concentration at 24h after dosing from one of three experiments in which cows were dosed experimentally with diclofenac. We chose to use the experiment which gave the highest ratio (Experiment 1). (3) We used the resulting ratio to multiply the mean concentration of tolfenamic acid in livers of experimentally treated cattle sampled 24h after dosing. Hence, our MLE dose is for a vulture which eats a large meal composed exclusively of liver from a treated animal. Concentrations of tolfenamic acid in other tissues, such as muscle, are likely to be much lower (EMEA 1997). This combination of choices is therefore highly precautionary. For example, had we decided to adopt the 90^th^ percentile in item (1) and the median result from item (2), by using the result from Experiment 2, the MLE dose in our experiments would have been one-third of that we used. It may be that a small proportion of individual vultures are more susceptible than others to a high dose of tolfenamic acid, as has been shown to be the case for *Gyps* vultures dosed with the NSAID ketoprofen (Naidoo et al. 2010). In future, we intend to compare the results of our MLE calculation with direct measurements of the concentration of tolfenamic acid in samples of liver and other tissues taken from cattle at carcass dumps in India, which represent the concentrations of the drug actually present in the food supply of wild vultures. However, such measurements are not yet available.

Our conclusion that tolfenamic acid is likely to be safe to wild vultures is supported by the fact that, not only did the remaining birds survive, but there was little evidence of an effect on the various blood serum parameters measured. Importantly, although blood serum concentrations of uric acid tended to increase after treatment, there was only a maximum of a doubling of concentrations in the 95% of birds that survived treatment, which is similar to that reported for meloxicam (Swan *et al.* 2006, Swarup *et al.* 2007). This compares with the 20+-fold increase in the birds that died after treatment with tolfenamic acid, and the similar increases reported for diclofenac and ketoprofen which are certainly nephrotoxic to *Gyps* vultures (Oaks *et al.* 2004, Swan *et al.* 2006, Naidoo *et al.* 2010).

Tolfenamic acid is increasingly being used to treat injured cattle in the subcontinent (Galligan *et al.* 2020), especially in Nepal and Bangladesh, and is manufactured in India and

Bangladesh, so promoting it in addition to meloxicam as a second vulture-safe drug would be feasible. The drug is intended for use in cattle as an anti-inflammatory, analgesic and anti-pyretic (Sidhu *et al.* 2010); it is also used in conjunction with antibiotics in the treatment of respiratory disease (Deleforge *et al.* 1994). In experiments comparing its efficacy to that of meloxicam in the reduction of stress and pain associated with the surgical castration of piglets, tolfenamic acid tended to be the more efficient analgesic (Wavreille *et al.* 2012). There is also evidence that it can help improve conception rate in cattle (Singh *et al.* 2020).

## CRediT authorship contribution statement

**Chandra Mohan:** Formal analysis, Investigation, Data Curation, Writing – Review & Editing. **A.K. Sharma:** Methodology, Formal Analysis, Investigation. **John Mallord:** Writing – Original Draft, Project administration. **Krishna Chutia:** Formal analysis, Investigation. **Reena Gupta:** Formal analysis, Investigation. **K. Mahendran:** Formal analysis, Investigation. **Kesavan Manickam:** Formal analysis, Investigation. **Karikalan Mathes:** Formal analysis, Investigation. **Abhijit Pawde:** Formal analysis, Investigation. **Nikita Prakash:** Resources, Project administration. **P. Ravichandran:** Formal analysis, Investigation. **Debasish Saikia:** Formal analysis, Investigation. **Rohan Shringarpure:** Formal analysis, Investigation. **Avinash Timung:** Formal analysis, Investigation. **Toby Galligan:** Methodology, Project administration. **Rhys Green:** Conceptualization, Methodology, Writing – Review & Editing. **Vibhu Prakash:** Conceptualization, Methodology, Writing – Review & Editing.

## Acknowledgments

We are grateful to Eurofins Advinus Ltd, Bengalaru, India for extraction and quantification of tolfenamic acid in plasma samples, and BNHS and IVRI for their general support. The work was funded by the RSPB, the Ministry of Environment, Forest and Climate Change of the Government of India and the Haryana State Forest Department.

