## Supplementary Information for "Experimental safety testing shows that the NSAID tolfenamic acid is not toxic to *Gyps* vultures in India at concentrations likely to be encountered in cattle carcasses"

**Appendix 1**

**Estimating the Maximum Likely Exposure (MLE) of wild vultures to veterinary drugs in the tissues of domesticated ungulates: a new approach and a worked example for tolfenamic acid**

**Introduction**

Experimental tests of the safety of NSAIDs to captive *Gyps* vultures require an estimate to be made of the maximum concentration of the drug in tissues of domesticated ungulates likely to be encountered by wild vultures under field conditions; the likely Maximum Level of Exposure (MLE). Several approaches have been adopted to calculating MLEs for NSAIDs in Gyps vultures. Principally, they involve analysing data from the experimental administration of a standard veterinary dose of the drug to cattle, followed by the measurement of the concentration of the drug in various tissues of the treated animals in relation to time since last administration of the drug. Sometimes the peak concentration of the drug in the tissue with the highest concentration is used from a series of determinations closely spaced in time (Naidoo *et al.* 2010). Another approach is to first determine the time after the last administration at which the concentration of the drug in plasma is highest and then sample tissues to determine drug concentration at that time, again selecting the tissue with the highest concentration (Naidoo *et al.* 2017). The concentration of the drug in this tissue is then converted to the MLE dose for a captive vulture by calculating the total quantity of drug expected to occur in a large meal of the tissue, which is usually taken to be the amount of tissue required to meet a free-living vulture’s energetic requirements for three days (Swan *et al.* 2006).

Two problems with this approach are (1) that it assumes that a standard veterinary course of the drug is actually administered by livestock owners, vets and paravets to domesticated ungulates under field conditions in the vulture range state and (2) that it neglects variation in the dose administered, the metabolism of the drug in cattle and the time elapsed between drug administration and death of the cow. Problem (1) has sometimes been addressed by assuming that typical users administer twice the standard veterinary dose (Naidoo *et al.* 2010), but this is an arbitrary procedure. In this appendix, we describe a new approach in which the empirical probability distribution of concentrations of diclofenac measured in the livers of cattle at carcass dumps in India is compared with concentrations of the same drug at various times after the last dose, as measured in cattle given a standard dose under experimental conditions. We argue that the ratio of a percentile of the empirical carcass dump distribution to the mean concentration of diclofenac in experimental cattle at a given time after the last dose can be used to scale up the concentration of another drug, measured in experiments at the same time elapsed as for diclofenac, to give a reasonable precautionary value for the MLE for the second drug.

**Methods**

*Empirical distribution of diclofenac concentrations in livers of of cattle at carcass dumps in India*

We used results from part of a time series of determinations of liver concentrations of diclofenac reported previously by Cuthbert *et al.* (2014). We used assays for diclofenac on liver samples from all 2,614 domesticated ungulates sampled in India in the years 2008-2010. The sampling was done at ungulate carcass dumps in the states of Andhra Pradesh, Haryana, Jammu & Kashmir, Madhya Pradesh, Maharashtra, Punjab, Rajasthan, Uttar Pradesh and West Bengal. Details of sampling and chemical assays are given in Cuthbert *et al.* (2014). . Of these samples, 135 had quantifiable levels of diclofenac (>0.01 mg/kg wet weight). We plotted the exceedance (negative cumulative) distribution of these 135 values and determined the 90^th^ and 95^th^ percentiles of the empirical distribution.

*Concentration of diclofenac in livers of experimentally treated cattle*

We used the results of three experiments on European and Indian cattle reported by Green *et al.* (2006). Experiment 1 was conducted on female Indian cattle *Bos indicus*. Animals received one injection of 1.0 mg/kg live weight of diclofenac. Experiment 2 was conducted on 16 young European cattle *B. taurus*. Animals received injections of 2.5 mg/kg live weight of diclofenac on six consecutive days. Experiment 3 was conducted on 8 mature female European cattle. Animals received injections of 2.5 mg/kg live weight of diclofenac on six consecutive days. Details are given in Green *et al.* 2006). Green *et al.* reported the results of a regression analysis of log-transformed diclofenac concentrations in each of several tissues of the treated cattle, in which log-concentration was regressed on time elapsed since the last dose of diclofenac. Here we use results for liver from the regression model presented in Table 1 of Green *et al.* (2006). The raw data for that analysis are shown graphically in Figure 1 of that paper.

**Results**

*Comparison of the empirical distribution of diclofenac concentrations in livers of cattle at carcass dumps in India with mean concentrations in livers of experimentally treated cattle*

Figure S1 shows the biphasic decline in mean diclofenac concentration in liver from the three experiments on cattle, as reported previously by Green *et al.* (2006). At a given time elapsed since the last dose, liver concentrations were higher for Experiments 2 and 3 than for Experiment 1, as would be expected from the larger dose and number of doses given in Experiments 2 and 3 than in Experiment 1 (Figure S1). When the empirical distribution of diclofenac concentrations in livers of cattle sampled at carcass dumps in India was compared with the mean levels from the experiments, it was found that the 90^th^ and 95^th^ percentiles of the empirical distribution were substantially higher than the mean values from the experiments at most times elapsed since the administration of the last dose (Figure S1; Table S1).

*Application to the determination of MLE for an NSAID with an unknown empirical distribution of liver concentration in cattle at carcass dumps*

In the case of many NSAIDs, experimental determinations of concentration of the drug in the tissues of cattle has only been done at one or a few times elapsed after administration of the last dose to the experimental animals. Often these times are long after the last dose and therefore well after the time at which peak concentration in liver is likely to have occurred. An example of such a drug is tolfenamic acid, for which the European Agency for the Evaluation of Medicinal Products (EMEA) reported the results of dosing cattle by intramuscular injection with two doses of tolfenamic acid at 2.0 mg/kg liveweight 48 hours apart (EMEA 1997). The mean concentration of tolfenamic acid in the liver 24h after the last injection was 1.4 mg/kg. Four days after the last treatment, tolfenamic acid was not present at quantifiable levels in the liver, so there was only a single usable value, which is long after the time at which peak concentrations of the drug in tissues are likely to have occurred. To obtain an MLE value for tolfenamic acid from these limited data, we first calculated the ratios of the 90^th^ and 95^th^ percentiles of the empirical distribution for diclofenac to the mean diclofenac concentration in experimental cattle at 24h after the last dose. We argue that a reasonable value for the MLE for tolfenamic acid can be obtained by multiplying the mean concentration of tolfenamic acid at 24 h post-treatment from the EMEA experiment by one of these ratios. Which percentile to use is arbitrary, so we give a range of values for the ratios, in the range 3 to 14, dependent upon which diclofenac experiment was used and which percentile of the empirical distribution was selected (Table S1). It can be argued that a precautionary approach is to use the highest of the ratios presented (13.9). In that case, the total quantity of tolfenamic acid in a wild vulture exposed to the MLE would be that present in a large meal (that taken if the bird was feeding at three-day intervals) of liver containing 13.9 times the mean 24 h concentration in the EMEA experiment: that is, 13.9 multiplied by 1.4 mg/kg = 19.5 mg per kg of liver. There may be justification for choosing other lower values of the ratio from Table S1, but we suggest that this seems to be a suitably precautionary approach.

| **Table S1.** 90^th^ and 95^th^ percentiles of the distribution of liver diclofenac concentrations for135 domesticated ungulates (mostly cattle and water buffaloes) with detectable diclofenac from surveys of carcass dumps in India in 2008 – 2010 and mean concentrations of diclofenac in the livers of experimentally treated cattle at 24h after the last dose was administered. Ratios of the carcass dump percentiles to the 24h experimental means are shown in the lower part of the table. | | |
| --- | --- | --- |
| *Percentile of carcass dump ungulate liver distribution* | | |
|  | Concentration (mg/kg) |  |
| 90th | 2.09 |  |
| 95th | 3.25 |  |
| *Mean concentration in cow liver at 24 h after administration of the last dose from experiments* | | |
|  | Concentration (mg/kg) |  |
| Experiment 1 | 0.23 |  |
| Experiment 2 | 0.47 |  |
| Experiment 3 | 0.70 |  |
| *Ratios of liver percentile concentration from carcass dump survey to experimental mean liver concentration at 24 h* | | |
|  | 90th percentile | 95th percentile |
| Experiment 1 | 8.9 | 13.9 |
| Experiment 2 | 4.5 | 6.9 |
| Experiment 3 | 3.0 | 4.6 |

**Figure S1.** Mean concentration of diclofenac in the livers of cattle in relation to time elapsed since the administration of the last dose from three experimental studies (Experiments 1, 2 and 3 of Green et al. 2006). These results are shown by the red, green and blue solid lines and the left-hand vertical axis. Also shown (stepped gold line) is the exceedance distribution (negative cumulative distribution) of diclofenac in the livers of all 135 domesticated ungulates (mostly cattle and water buffaloes) with detectable diclofenac from surveys of carcass dumps in India in 2008 – 2010. The exceedance probability is the proportion of samples (on the right-hand vertical axis) with a concentration at or in excess of the value shown on the horizontal axis. The vertical gold lines show the 90th and 95th percentiles of the distribution of values for animals with detectable diclofenac. It can be seen that the 90th and 95th percentiles of the distribution are much larger (from 3 to 14 times) than the mean liver concentrations of diclofenac in experimental cattle at 24 h after the last dose, as shown by the vertical red, green and blue dashed lines.

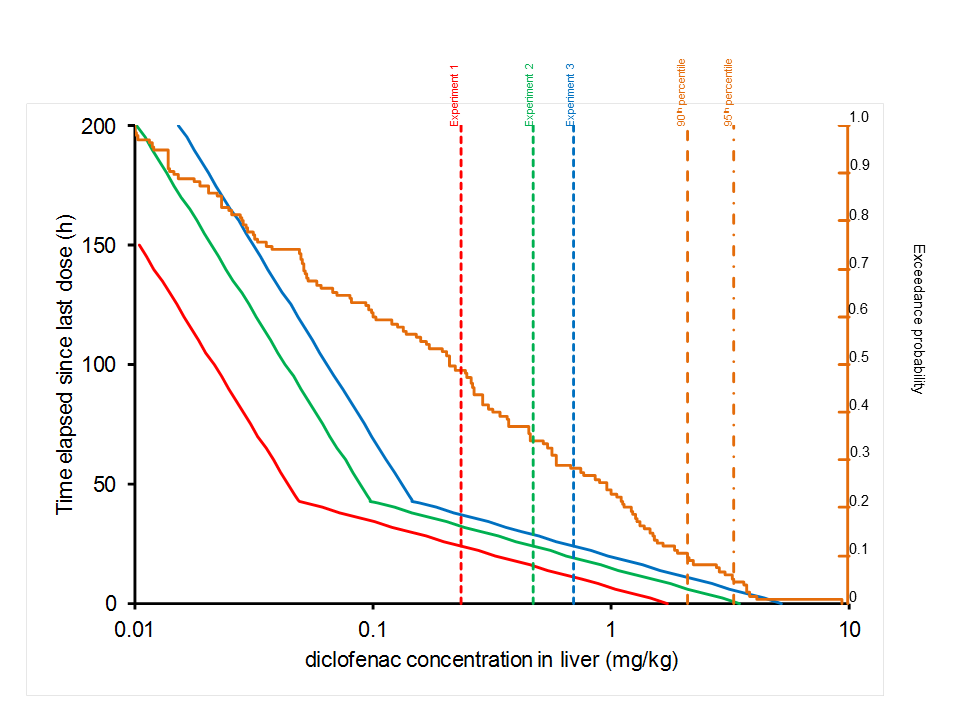

**Appendix 2**

**Concentrations of constituents of blood serum of Himalayan Griffon vultures in relation to time since dosing with tolfenamic acid**

**Table S2.** Concentrations (mg l^-1^) of constituents of blood serum of Himalayan Griffon vultures in relation to time since dosing with tolfenamic acid by oral gavage.

|  | Albumin |  |  |  | Total protein |  |  |  | Phosphorus |  |  |  | Urea |  |  |  |
| --- | --- | --- | --- | --- | --- | --- | --- | --- | --- | --- | --- | --- | --- | --- | --- | --- |
|  | Control | Dosed |  |  | Control | Dosed |  |  | Control | Dosed |  |  | Control | Dosed |  |  |
| Time h | Mean (n) | Mean (n) | *t* | *p* | Mean (n) | Mean (n) | *t* | *p* | Mean (n) | Mean (n) | *t* | *p* | Mean (n) | Mean (n) | *t* | *p* |
| 0 | 8.7 (15) | 10.3 (36) | 1.38 | 0.17 | **2.8733 (15)** | **3.192 (36)** | **2.17** | **0.035** | 1.7667 (9) | 2.0586 (29) | 0.72 | 0.48 | 7.85 (11) | 6.86 (31) | 0.55 | 0.59 |
| **2** | 7.5 (15) | 7.9 (36) | 0.75 | 0.46 | 2.9867 (15) | 3.139 (36) | 0.97 | 0.34 | 2.4333 (7) | 2.5222 (29) | 0.19 | 0.85 | 7.92 (11) | 10.56 (29) | 1.52 | 0.14 |
| **6** | **6.9 (15)** | **8.0 (36)** | **2.45** | **0.02** | 2.86 (15) | 3.142(36) | 2.0 | 0.051 | 2.3555 (9) | 2.4207 (29) | 0.13 | 0.90 | 7.51 (11) | 8.81 (31) | 0.52 | 0.61 |
| 12 | 8.3 (15) | 9.7 (36) | 1.39 | 0.17 | 3.4 (15) | 3.639 (36) | 0.86 | 0.40 | 2.3333 (9) | 2.4586 (29) | 0.28 | 0.78 | **6.56 (11)** | **10.29 (31)** | **2.58** | **0.014** |
| 24 | 8.7 (15) | 9.8 (36) | 1.31 | 0.20 | 3.0267 (15) | 3.139 (36) | 0.57 | 0.57 | 2.35 (6) | 2.6158 (19) | 0.46 | 0.65 | 6.1 (7) | 7.79 (14) | 0.79 | 0.44 |
| 36 | 8.7 (15) | 9.6 (36) | 1.08 | 0.29 | **2.8267(15)** | **3.2(36)** | **2.25** | **0.029** | 2.25 (6) | 2.5765 (17) | 0.67 | 0.51 | **5.81 (7)** | **14.53 (12)** | **2.82** | **0.012** |
| 48 | **6.7 (15)** | **8.3 (36)** | **2.92** | **0.005** | 3.0333 (15) | 3.164 (36) | 0.59 | 0.56 | 2.1833 (6) | 2.4294 (17) | 0.6 | 0.56 | 6.64 (7) | 7.59 (14) | 0.32 | 0.75 |
| 96 | 8.5 (11) | 10.5 (19) | 1.92 | 0.06 | 3.0818 (11) | 3.247 (19) | 0.56 | 0.58 | - | - | - | - | 8.42 (7) | 14.88 (14) | 2.0 | 0.061 |
| 168 | 8.3 (8) | 7.4 (9) | 1.07 | 0.30 | 3.225 (8) | 3.4889 (9) | 1.04 | 0.31 | - | - | - | - | - | - | - | - |
|  |  |  |  |  |  |  |  |  |  |  |  |  | Tolfenamic |  |  |  |
|  | Creatinase |  |  |  | Calcium |  |  |  | Chlorine |  |  |  | acid |  |  |  |
|  | Control | Dosed |  |  | Control | Dosed |  |  | Control | Dosed |  |  | Survived | Died |  |  |
| Time h | Mean (n) | Mean (n) | *t* | *p* | Mean (n) | Mean (n) | *t* | *p* | Mean (n) | Mean (n) | *t* | *p* | Mean (36) | Mean (2) | *t* | *p* |
| 0 | 0.718 (11) | 0.777 (31) | 0.49 | 0.62 | 9.14 (15) | 9.197 (36) | 0.11 | 0.92 | 107.97 (11) | 117.25 (31) | 1.25 | 0.22 | - | - |  |  |
| **2** | 0.836 (11) | 0.797 (29) | 0.31 | 0.76 | 8.807 (15) | 8.436 (36) | 0.76 | 0.45 | 105.47 (11) | 114.81 (29) | 1.28 | 0.21 | 5647.895 | 5935 | 0.13 | 0.90 |
| 6 | 0.982 (11) | 1.052 (31) | 0.70 | 0.49 | 8.813 (15) | 9.403 (36) | 1.22 | 0.23 | 106.05 (11) | 105.22 (31) | 0.12 | 0.90 | 2258.632 | 2005.5 | 0.29 | 0.77 |
| 12 | 0.864 (11) | 0.868 (31) | 0.05 | 0.96 | **7.767 (15)** | **8.894(36)** | **2.26** | **0.029** | 104.16 (11) | 112.24 (31) | 1.50 | 0.14 | 1033.95 | 625.5 | 0.82 | 0.42 |
| 24 | 1.009 (11) | 1.039 (31) | 0.23 | 0.82 | 8.307 (15) | 8.353 (36) | 0.12 | 0.90 | 108.97 (11) | 122.64 (31) | 1.84 | 0.07 | 291.870 | 167 | 0.55 | 0.58 |
| 36 | 1.009 (11) | 1.162 (29) | 1.31 | 0.20 | 8.58 (15) | 8.442 (36) | 0.31 | 0.76 | 110.95 (11) | 119.0 (29) | 1.37 | 0.18 | 131.151 | 147 | 0.17 | 0.86 |
| 48 | 1.073 (11) | 0.952 (31) | 1.11 | 0.27 | 8.08 (15) | 9.086 (36) | 1.73 | 0.09 | 106.93 (11) | 109.45 (31) | 0.54 | 0.59 | 49.903 | - | - | - |
| 96 | 0.936 (11) | 0.843 (14) | 0.76 | 0.46 | 10.03 (11) | 9.037 (19) | 1.48 | 0.15 | 110.4 (11) | 105.9 (14) | 0.98 | 0.34 | 6.8892 | - | - | - |
| 168 | 1.025 (4) | 0.975 (4) | 0.28 | 0.79 | 8.613 (8) | 8.5111 (9) | 0.17 | 0.87 | 113.83 (4) | 115.28 (4) | 0.29 | 0.78 | 0.570 | - | - | - |

**Table S3.** Concentrations (mg l^-1^) of constituents of blood serum of Himalayan Griffon vultures (n = 4) in relation to time since dosing with tolfenamic acid after feeding with contaminated buffalo meat, compared to control group (n = 2).

|  | Albumin |  |  |  | Total protein |  |  |  | Phosphorus |  |  |  | Urea |  |  |  |
| --- | --- | --- | --- | --- | --- | --- | --- | --- | --- | --- | --- | --- | --- | --- | --- | --- |
|  | Control | Dosed |  |  | Control | Dosed |  |  | Control | Dosed |  |  | Control | Dosed |  |  |
| Time h | Mean | Mean | *t* | *p* | Mean | Mean | *t* | *p* | Mean | Mean | *t* | *p* | Mean | Mean | *t* | *p* |
| 0 | **5.7** | **6.7** | **3.54** | **0.02** | 2.3 | 2.5 | 1.65 | 0.17 | 1.4 | 2.5 | 1.37 | 0.24 | 9.6 | 9.3 | 0.36 | 0.74 |
| 48 | 6.6 | 6.0 | 0.65 | 0.55 | 2.6 | 2.3 | 0.84 | 0.45 | 1.8 | 2.7 | 1.24 | 0.28 | 9.0 | 14.1 | 1.01 | 0.37 |
| 168 | 6.6 | 6.5 | 0.14 | 0.90 | 2.6 | 3.0 | 1.02 | 0.36 | 2.0 | 1.9 | 0.28 | 0.80 | 8.8 | 11.1 | 2.04 | 0.11 |
|  |  |  |  |  |  |  |  |  |  |  |  |  | Tolfenamic |  |  |  |
|  | Creatinase |  |  |  | Calcium |  |  |  | Chlorine |  |  |  | acid |  |  |  |
|  | Control | Dosed |  |  | Control | Dosed |  |  | Control | Dosed |  |  | Control | Died |  |  |
| Time h | Mean (n) | Mean | *t* | *p* | Mean | Mean | *t* | *p* | Mean | Mean | *t* | *p* | Mean | Mean | *t* | *p* |
| 0 | 0.7 | 1.5 | 1.29 | 0.27 | 8.5 | 9.1 | 0.31 | 0.77 | 117.3 | 114.9 | 1.83 | 0.14 | 0 | 0 | - | - |
| 48 | 0.7 | 0.8 | 2.31 | 0.08 | 7.6 | 10.5 | 1.27 | 0.27 | 106.7 | 110.3 | 0.80 | 0.47 | 2.8 | 13.2 | - | - |
| 168 | 1.0 | 0.9 | 0.84 | 0.45 | 7.7 | 7.9 | 0.13 | 0.90 | 119.9 | 116.5 | 0.99 | 0.38 | 0 | 0 | - | - |
